# Atypical development of subcortico-cortical effective connectivity in autism

**DOI:** 10.1101/2021.07.02.450977

**Authors:** Luigi Lorenzini, Guido van Wingen, Leonardo Cerliani

## Abstract

Hypersensitivity, stereotyped behaviors and attentional problems in autism spectrum disorder (ASD) are compatible with inefficient filtering of undesired or irrelevant sensory information at early stages of neural processing. This could stem from delays in the neurotypical development of the functional segregation between cortical and subcortical brain processes, as suggested by previous findings of overconnectivity between primary sensory regions and deep brain nuclei in ASD.

To test this hypothesis, we used dynamic causal modelling to quantify the effect of age on the development of (1) cortical functional segregation from subcortical activity and (2) directional influence of subcortical activity on cortical processing in 166 participants with ASD and 193 typically developing controls (TD) from the Autism Brain Imaging Data Exchange (ABIDE).

We found that in TD participants age was significantly associated with increased functional segregation of cortical sensory processing from subcortical activity, paralleled by a decreased influence of subcortical activity on cortical processing. Instead these effects were highly reduced and mostly absent in ASD participants, suggesting a delayed or arrested development of the segregation between subcortical and cortical sensory processing in ASD.

This atypical configuration of subcortico-cortical connectivity in ASD can result in an excessive amount of unprocessed sensory information relayed to the cortex, which is likely to impact cognitive functioning in everyday situations where it is beneficial to limit the influence of basic sensory information on cognitive processing, such as activities requiring focused attention or social interactions.

## 1. Introduction

During the transition from childhood to adulthood our brain develops the ability to determine behavior on the basis of abstract representations and long-term plans, which are not primarily driven by current sensory stimuli, emotions and interoceptive feelings. This transition is reflected in the development of brain connectivity, and specifically in the increased independence of cortical information processing from subcortical inputs, coupled with the strengthening of long-range cortico-cortical connections within and between large-scale brain networks supporting higher-order distributed cognitive functions (K. Supekar, Musen, and Menon 2009).

In autism, the development of brain connectivity follows an atypical trajectory, and appears to be delayed or arrested at an immature stage. This is suggested by the underconnectivity between posterior and anterior brain regions within default mode, attentional and language networks (Herbert et al. 2003; Just et al. 2004; Muller et al. 2011; Kana et al. 2014), together with the persistent overconnectivity between cortical, subcortical and cerebellar regions (Di Martino et al. 2011; Cerliani et al. 2015; Woodward et al. 2017; Oldehinkel et al. 2019; Maximo and Kana 2019). Such a connectivity pattern hampers the development of functional segregation between cortical networks (Holiga et al. 2019; Müller and Fishman 2018), leading to an atypical integration of information among them (Hong et al. 2019; Jeffrey D. Rudie et al. 2012; Fishman et al. 2014). Specifically, recent neuroimaging studies suggest that functional integration in ASD is highly driven by current sensory information. In this respect, Hong and colleagues (Hong et al. 2019) showed that in autism spectrum disorders (ASD) the information encoded in primary sensory cortices flows more rapidly to transmodal regions. This situation increases the amount of basic sensory information reaching attentional and associative networks, and therefore the relevance of current sensory stimuli in determining behavior. A similar conclusion can be drawn from another study (Holiga et al. 2019) where decreased intrinsic functional connectivity of sensory and higher order fronto-parietal networks in ASD was associated with increased cross-talk between them. Finally, the presence of overconnectivity between subcortical and primary sensory regions (Cerliani et al. 2015; Woodward et al. 2017; Maximo and Kana 2019) suggests that deficits in filtering unwanted or irrelevant sensory stimuli in ASD might originate in early stages of sensory input processing, at the subcortical level.

The defective functional segregation of primary sensory regions, as well as the abnormally high influence of subcortical regions over cortical processing, likely reflects the atypical development of brain connectivity in ASD, as several studies reported an age-related decrease in subcortico-cortical functional connectivity in typically developing participants but not in ASD (Iidaka et al. 2019; Cerliani et al. 2015). However these previous studies could not directly investigate the functional segregation of primary sensory regions and the directional, bottom-up influence of subcortical over cortical regions, since the results were based on symmetric measures of functional connectivity - for instance the Pearson correlation coefficient - which do not yield a causal interpretation of brain dynamics, known as effective brain connectivity (Karl J. Friston 2011) (differences between functional and effective connectivity are illustrated in Figure 1). Therefore, in the present study we used spectral dynamic causal modelling (DCM, (K. J. Friston, Harrison, and Penny 2003)) to investigate the interaction between subcortical nuclei and primary sensory cortical regions in resting-state fMRI data (Karl J. Friston et al. 2014). Specifically, we examined (1) the directional influence of subcortical activity on cortical sensory processing and (2) the intrinsic inhibition of each region, which reflects its sensitivity to the influence of other brain regions in the model (i.e. functional segregation) in both ASD and TD participants. We hypothesized age to be associated with a decrease in bottom-up connectivity and increased functional segregation of cortical regions in both ASD and TD participants. Crucially, however, we hypothesized that this age-dependent effect would be significantly attenuated in ASD with respect to TD participants.

**Figure 1.**
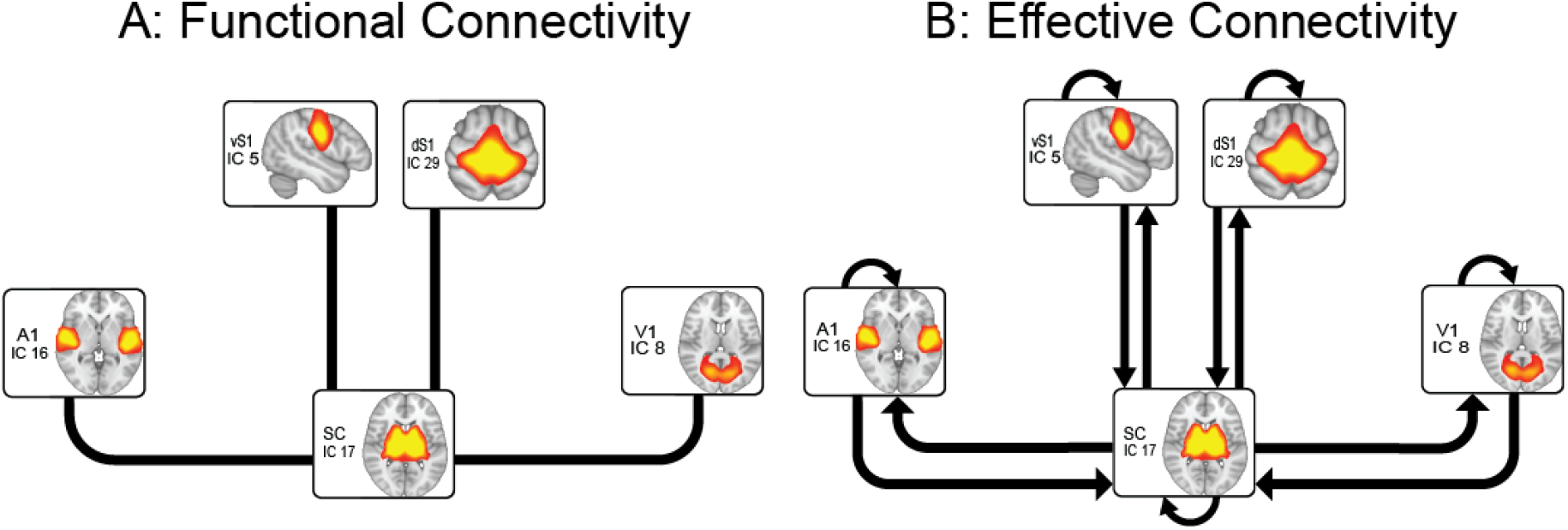
Functional and effective connectivity. **A:** Functional connectivity captures patterns of statistical dependence between regions of interest (ROIs) through the correlation of their fMRI time-series. Five ROIs, including subcortical nuclei (basal ganglia and thalamus) and primary sensory regions (dorsal and ventral somatosensory, primary auditory and primary visual cortex) showed increased functional connectivity in ASD compared to TD in our previous work (Cerliani et al. 2015). **B:** Effective connectivity models causal influences that one neural network exerts onto another. The figure depicts the connections we chose to model in our DCM analysis: *bottom-up* influence of subcortical nuclei on the primary sensory cortices; *top-down* influence of primary sensory regions on subcortical activity; and *auto-connections*, which in DCM model inhibitory self-connection of one neural system with itself and reflect its functional segregation - that is the sensitivity of a region to the influence of another modelled input (Zeidman, Jafarian, Corbin, et al. 2019).

## 2. Methods

### 2.1 Participants

We included in our study 166 participants with high-functioning ASD (all male, median age = 17.6, sd = 7.6) and 193 typically developing controls (TD - all male, median age = 16.9, sd = 6.6) sampled from the 1111 dataset in the Autism Brain Imaging Data Exchange dataset (ABIDE I - A. Di Martino et al. 2013). The selection procedure (detailed in Figure S1) ensured that (1) ASD and TD participants were matched by age, IQ, head motion and eye status in the scanner at the group level, (2) the images were devoid of problematic artifacts arising from image acquisition issues or motion. Table 1 reports the demographics of the final sample of 359 participants. Supplementary figures S4 and S5 provide additional information on the distribution of participants’ age per site and across sites.

**Table 1.** Participants Demographics and Cognitive Scores. Abbreviations: ASD, autism spectrum disorder group; TD, typically development group; N/A, not applicable; ADI-R, Autism Diagnostic Interview - Revised (C. Lord, Rutter, and Le Couteur 1994); ADOS, Autism Diagnostic Observation Schedule (Catherine Lord et al. 1989); SRS, Social Responsiveness Scale (Constantino et al. 2003; Constantino and Todd 2003).

|  | Mean (SD) [Range] |
| --- | --- |

|  | ASD (N = 166) | TD (N = 193) |
| --- | --- | --- |
| Age, years | 17.6 (7.6) [7-50] | 16.9 (6.6) [6.5-39.4] |
| Full-scale IQ | 109.6 (16.16) 71-148 | 111 (13.1) [73-146] |
| ADI-R Social (N=93) | 19.7(5.3)[7-28] | N/A |
| ADI-R Verbal (N=94) | 15.6(4.5)[2-25] | N/A |
| ADI-R Repetitive Behavior (N=93) | 5.8(2.58)[0-12] | N/A |
| ADOS Total (N=171) | 10.7(5.3)[0-22] | N/A |
| ADOS Communication (N=170) | 3.5(1.9)[0-8] | N/A |
| ADOS Social (N=171) | 7.1(3.8)[0-14] | N/A |
| ADOS Repetitive behavior (N=142) | 1.7(1.6)[0-8] | N/A |
| SRS (N<sub>ASD</sub> = 111; N<sub>TD</sub> = 108) | 89.4 (32.4) [6-164] | 22.2 (18.1) [0-103] |

### 2.2 Selection of the regions of interest for the analysis of effective connectivity

We aimed to estimate the effective connectivity between subcortical nuclei and primary sensory regions during resting-state fMRI. To determine the location of these regions in a data-driven way, we carried out an independent component analysis (ICA) using FSL Melodic meta-ICA (Beckmann and Smith 2004). All the details about data preprocessing and meta-ICA are described in detail in a previous study (Cerliani et al. 2015) and in the supplementary materials. Notably, we strived to remove motion by (1) regressing the estimated motion parameters (2) carrying out ICA Aroma (Pruim et al. 2015) and (3) excluding participants featuring relatively high residual motion quantified by framewise displacement (mean framewise displacement across all time points > 0.34). Implementing meta-ICA (Biswal et al. 2010) allowed us to extract 19 spatially independent components mostly located in the gray matter, featuring high reproducibility across twenty-five subsets of participants and with high resemblance to functional networks recruited by task-based fMRI experiments (Smith et al. 2009; Laird et al. 2013). Our previous functional network connectivity analysis evidenced that among all components, the group of ASD participants showed a significantly higher interaction between one subcortical component - encompassing the basal ganglia and thalamus - and four primary sensory cortical networks - ventral and dorsal somatosensory, visual and auditory (Cerliani et al. 2015). Therefore in the present study we use the spatial maps associated with these 5 components (thresholded at Z > 3 from the meta-ICA results) to estimate differences in their functional segregation and directional interaction between ASD and TD participants using dynamic causal modelling (Karl J. Friston et al. 2014).

### 2.3 Spectral dynamic causal modelling

Dynamic Causal Modelling (DCM, (K. J. Friston, Harrison, and Penny 2003)) aims to model the effective connectivity - that is the causal interactions between brain regions - in order to derive information about the direction and strength of each connection. While traditionally DCM was applied only to task-based fMRI, recently spectral dynamic causal modelling (spDCM) was specifically devised to estimate effective connectivity in resting-state fMRI data (Karl J. Friston et al. 2014). spDCM uses a Bayesian framework to model directional interactions amongst brain regions based on their cross spectral densities and obtain estimates of the strength of each connection. Technical details on spDCM can be found in the Supplementary materials and in the reference papers (Razi et al. 2015, 2017).

Spectral DCM was carried out using SPM 12 (https://www.fil.ion.ucl.ac.uk/spm) (Karl J. Friston et al. 2007). Following well established procedures (Karl J. Friston et al. 2016), in the *first-level* (single subject) analysis we only specified one full model per subject, including all the bottom-up (subcortico-cortical) and top down (cortico-subcortical) connections between our regions of interest, as well as the inhibitory self-connections within each region (Fig 1B). Since direct connections between primary sensory cortices are anatomically implausible (Mesulam 2000), cortico-cortical connections were not modelled, reducing the number of parameters to be estimated. Inversion (fitting) of the model to the data provided an estimation of single-subject DCM parameters, i.e. connection strengths. Comparison with reduced models was carried in the *second-level* (group) analysis (see below). Explained variance of full DCM models was inspected to ensure convergence. No subject was excluded due to poor data fit.

### 2.4 Parametric empirical Bayes

To test differences in DCM parameters at the group level, we used a recent implementation of SPM to model group effective connectivity in the context of DCM, known as Parametric Empirical Bayes (PEB) (Karl J. Friston et al. 2016). In brief, this can be considered as a Bayesian second-level general linear model testing how subject measures (individual connection strengths) relate to the group mean and other group-level variables. This routine has the advantage of taking into consideration the full posterior density from the first level (single-subject) DCM to inform the second level results (Zhou et al. 2018).

The main PEB model included the mean, age, group and group-by-age interaction as between-subject variables of interest. To exemplify significant effects found in the interaction term, we specified a second model including the effect of age separately within the two groups. In each model, the rs-fMRI mean (across time points) framewise displacement (FD, (Power et al. 2012)) of each subject was also included as a nuisance variable, to model potential residual effects of movement. Finally, since the data of this ABIDE sample was collected in different sites, we also modelled site with additional dummy covariates (more on this below).

Following current standards (Karl J. Friston et al. 2016), Bayesian model reduction (BMR) was then used to estimate several nested (reduced) models by assuming one or more connections from the full model to be selectively switched off, and derived evidence directly from the full model. Bayesian model average (BMA) was subsequently employed to estimate a weighted average of the parameter strength based on nested models’ log evidence and estimate the influence of between subjects regressors. Further details on BMR and BMA procedures can be found in the supplementary materials and Figure S3.

### 2.5 Modelling site-related confounds

Data from the present ABIDE sample was acquired in 8 different sites. In order to model out this potential confound, we followed the standard procedure of introducing 7 dummy covariates (one less than the number of sites, to prevent rank deficiency of the model matrix) encoding each site as 1 for participants from that site, and 0 everywhere else. However, a preliminary analysis of variance revealed significant mean age difference across sites (F(7,351) = 22.07, p < 0.001, see Figure S5). Therefore, such dummy variables prevent the age predictor from capturing the variance which is shared between age and site-related confounds, as in the general linear model only the variance in the dependent variable which is unique to a particular predictor is captured by that predictor (Poldrack, Mumford, and Nichols 2011). In other words introducing dummy variables to control for the effect of site - given the significant differences in age across sites - introduces (partial) collinearity in the model and effectively removes variability in DCM estimates (the mean for each site) which is explained by both age and the dummy variables used to model inter-sites differences, rather than by confounding differences between sites only. Such procedure can potentially make the results unstable and introduce false negatives in the estimation of the association between age and effective connectivity, which represents the main scope of our analyses. For these reasons, we will report the results both with and without correction for site, since it is not possible, given the data at hand, to separately model the variance in connectivity estimates which is due to either site-related confounds or to interesting differences in the age of the participants.

### 2.6 Interpreting PEB results

In the context of DCM, the probability of the effects predicted by the model is estimated in a Bayesian framework. Bayesian statistics incorporates prior knowledge of the event in the model to be tested - for instance the presence of a connection between two brain areas. Contrary to the frequentist approach, which does not explicitly test the probability of the hypothesized effect, in the Bayesian framework we test the probability of our specific hypothesis given the observed data. Therefore, the frequentist concept of statistical significance does not apply to Bayesian statistics which in turn provides information on the likelihood of the hypothesized effect. Here, the PEB outcome provides two types of information: 1) the estimated effect, which refers to the strength and the sign of the influence that each covariate exerts on each DCM connection, expressed in Hz: for example, an effect of age on the self-connection of S1 of +0.14 means that the self-inhibition of S1 increases 0.14 times the age score; 2) the probability of the parameter, which represents the probability of the observed effect. Following previous DCM studies (Almgren et al. 2018), we will consider significant those experimental effects where the posterior probability of the DCM parameter given the data (P(M|Y)) exceeds 0.90, and therefore show a 90% confidence interval (hereafter CI) not including the zero (Makowski, Ben-Shachar, and Lüdecke 2019).

### 2.7 Association with symptoms severity

We then evaluated the relationship between effective connectivity and behavioural symptoms that are often observed in association with ASD, in a subsample of individuals for which Social Responsiveness Scale (SRS (Constantino et al. 2003)) scores are provided in the ABIDE dataset (n=219). Linear mixed models were used to study the effect of SRS and its interaction with age on estimated DCM parameters (connectivity strengths) of regions showing significant association in PEB analysis. As for the PEB model, FD was added as a covariate. A random intercept was used to correct for the effect of the site. P-values were adjusted for multiple comparisons using false discovery rate (FDR).

## 3. Results

In our previous investigation on this sample from the ABIDE dataset (Cerliani et al. 2015) we reported that resting-state functional connectivity between subcortical and primary sensory regions was increased in ASD with respect to TD participants (Fig. S2). In the present study we hypothesized that this subcortico-cortical overconnectivity could be explained by an atypical development of bottom-up projections and functional segregation of the cortex. To test this hypothesis we examined the relationship between age and DCM parameter estimates for functional segregation (self-inhibition) and directional bottom-up influences.

### 3.1 Main effect of Age across all participants

Figure 2 shows the main effect of age on bottom-up and top-down influence between subcortical and cortical sensory regions in the whole sample of 359 participants (ASD + TD). In addition, it shows the effect of age on self-connections, which model each region’s excitatory-inhibitory balance: a stronger inhibitory self-connection reflects higher functional segregation of that region’s activity from the influence of the other regions in the model (Zeidman, Jafarian, Seghier, et al. 2019).

**Figure 2.**
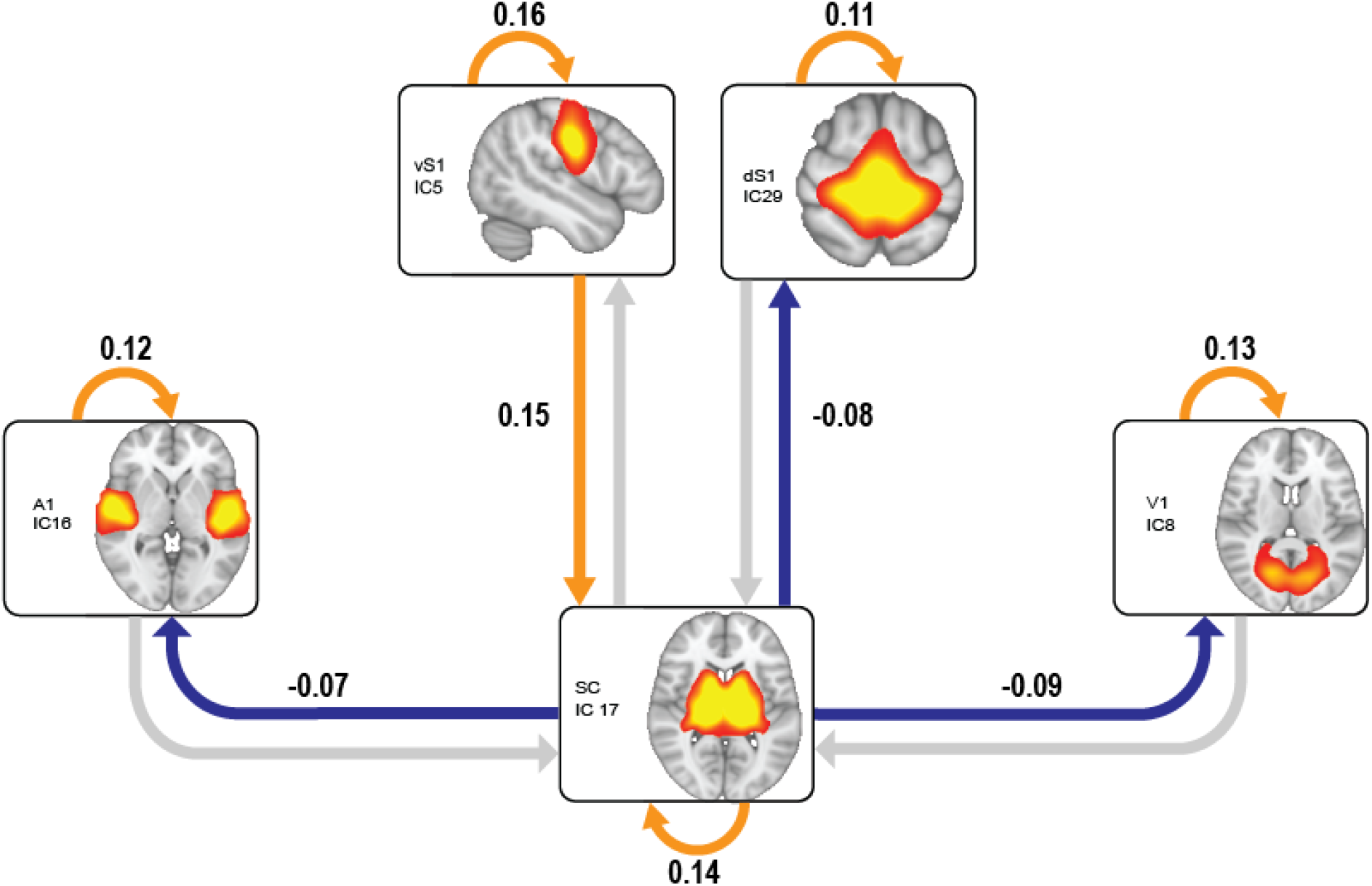
Main effect of Age. Association between Age and directed subcortico-cortical influence or regional functional segregation (self-connections) in the whole group. An increase in age results in decreased influence (in blue) of subcortical over cortical sensory activity, and increased segregation (in orange) of the intrinsic activity of each region from that of other regions in the model. In the case of the ventral somatosensory cortex, age is associated with an increased top-down influence (in orange) on subcortical activity. Orange and blue connections represent interactions showing a significant effect of age (90% CI on DCM parameter estimates outside zero). The DCM parameter estimate for each significant connection is reported close to the corresponding arrow. SPM graphical outputs and results of this analysis before and after site correction are reported in Figure 6 of supplementary materials.

Consistent with our predictions and previous literature, age was significantly associated with increasing functional segregation across all cortical and subcortical regions (self-connections in orange in Fig. 2). Age was also significantly associated with decreasing influence of subcortical activity on primary sensory regions across all sensory modalities (blue connections in Fig. 2). In the case of the ventral somatosensory cortex, age was significantly associated with an increasing top-down influence from the cortex on subcortical regions.

### 3.2 Reduced functional segregation of primary sensory regions in ASD vs TD

Figure 3A shows the DCM parameter estimates of functional segregation (self-connection) for each sensory region (and the associated 90% CI) when age was modelled separately in TD and ASD participants. The functional segregation of primary somatosensory (vS1, dS1) and auditory (A1) regions significantly increased with age in TD participants (black CI bars in Figure 3A) while in ASD participants this effect was significant only in the visual modality (V1). When these parameter estimates were compared between groups - in a single model including both ASD and TD - the age-by-group interaction confirmed that age has a smaller effect on the increase of cortical functional segregation in ASD in the somatosensory (vSI) and auditory modality (A1) (asterisks in Figure 3A).

**Figure 3.**
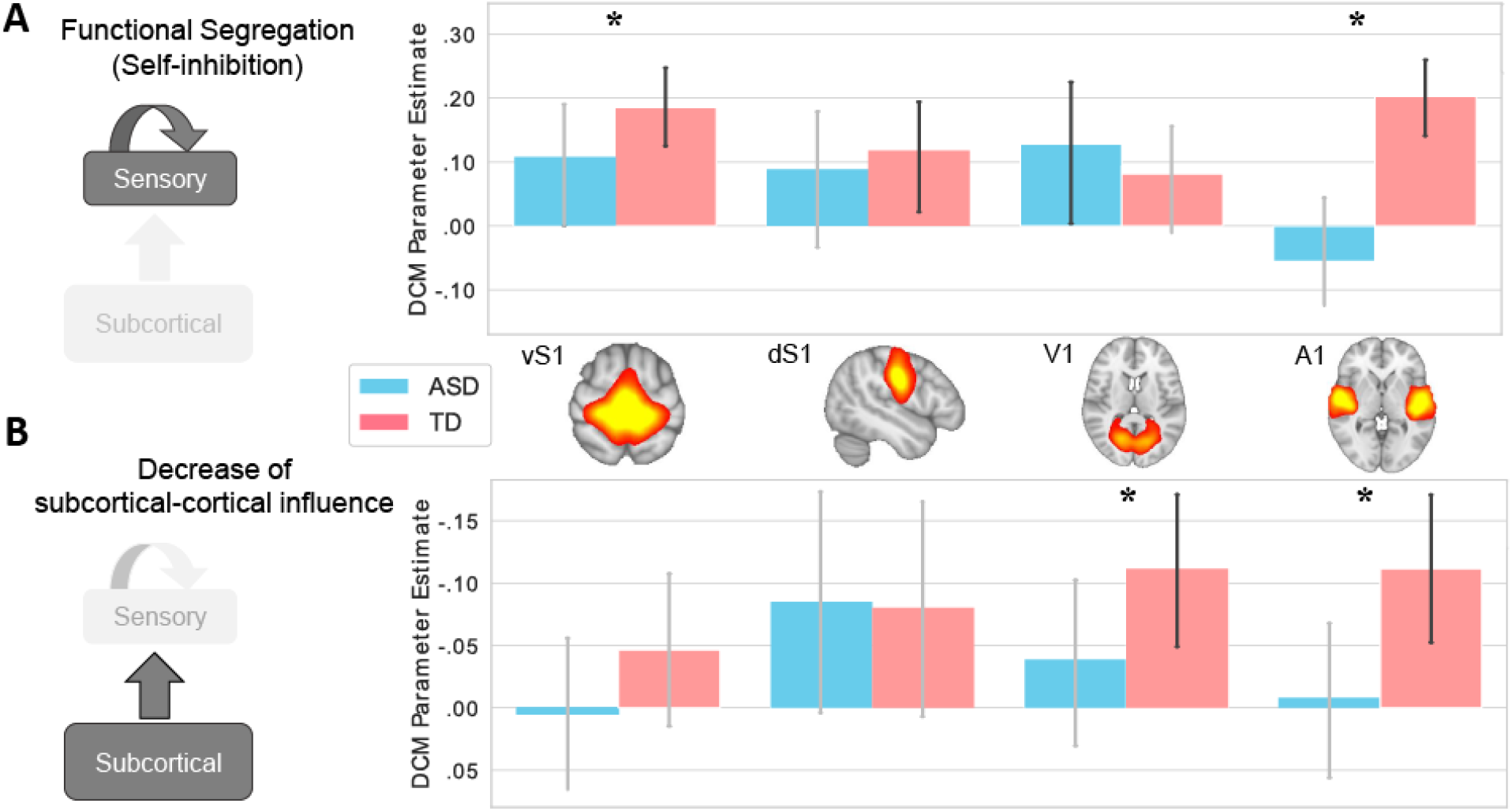
Age-related group differences in effective connectivity. (blue = ASD, pink = TD). Error bars indicate the 90% CI around the DCM parameter estimates for the within-group DCM, in which age was modelled separately for ASD and TD participants. In DCM, a 90% CI not including zero is used to determine if the parameter estimates are to be considered significant (Almgren et al. 2018). Black error bars reflect 90% CI not including zero, therefore indicating a significant effect of Age. Asterisks denote a significant (90% CI) age-by-group interaction (TD > ASD). **A: Effect of age on functional segregation.** In TD, age is significantly associated with an increase of the functional segregation of primary somatosensory (vS1, dS1) and auditory (A1) regions, while in ASD this is the case only for V1. The age-by-group interaction effect is significant in vS1 and A1. **B: Effect of age on subcortico-cortical influence:** In TD, age is significantly associated with a reduced influence of subcortical regions on cortical sensory processing in the visual (V1) and auditory (A1) modalities (note that the values on the Y axis are inverted). By contrast, age is not significantly associated with changes of bottom-up influence in ASD participants. The age-by-group interaction confirmed the presence of a significant difference between ASD and TD in the reduction of bottom-up connectivity for these sensory modalities. SPM graphical outputs and results of this analysis before and after site correction are reported in Figure 7 of supplementary materials.

This result shows that some primary sensory regions are more susceptible to be influenced by subcortical activity in ASD participants than in TD participants of comparable age. Conversely, age appeared to contribute to a higher functional segregation of the subcortical regions in ASD than in TD (not presented in Figure 3), but this effect only showed a trend towards significance (P(M|Y) = 0.88).

Importantly, these and the following results (next paragraph) were significant only in relation to the age of the participants, while no main effect of group was significant if age was treated as a confound.

### 3.3 Persistent subcortical influence on cortical sensory processing

Figure 3B shows the DCM parameter estimates for the influence of subcortical on primary sensory brain activity (and the associated 90% CI) when age was modelled separately in TD and ASD participants. Note that since we are showing the effect of age on the decrease in the influence of subcortical nuclei on primary sensory regions, we inverted the values on the Y axis. In TD participants, the subcortical influence on visual (V1) and auditory (A1) cortical activity significantly decreased with age, while this was not observed for any sensory modality in ASD participants. The age-by-group interaction confirmed that age was significantly associated with a stronger decrease of subcortical influence over visual and auditory primary cortical regions in TD than in ASD (Figure 3B: asterisks).

SPM graphical outputs of the described analysis are reported in the supplementary materials. Adding site into the PEB model did not change the direction of the effect and showed minimal reduction of posterior probabilities, though leading to non-significance for several nodes (Figure S6-S7).

### 3.4 Association with Social Responsiveness Scale

In order to examine the association between neuroimaging results and behaviour, we studied whether similar effective connectivity alterations could be observed in relation with total SRS score. The interaction between age and SRS showed a trend to significant effect only on auditory cortex self-connectivity (p-value = 0.053 after q(FDR)=0.05 correction). Specifically, the functional segregation of A1 decreased with age in participants with severe symptoms, while it remained stable or even increased in participants with mild or moderate behavioural symptoms (Figure 4). Models’ coefficients are shown in table S1 of supplementary materials.

**Figure 4.**
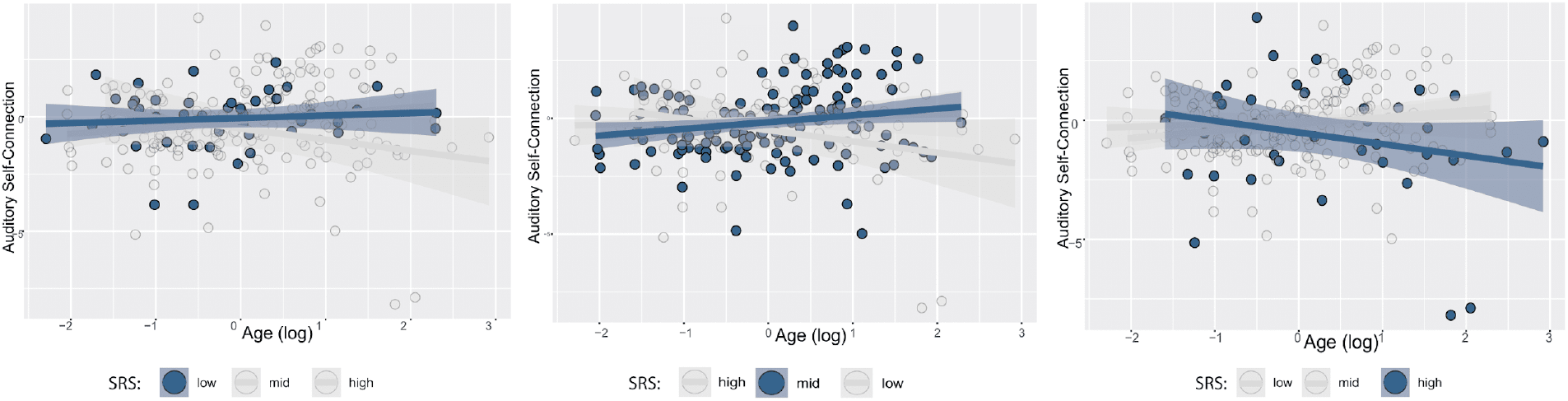
Effect of the interaction between age and SRS on DCM parameters. For clarity of presentation, participants’ SRS are stratified in low (total SRS < −1SD), mid (total SRS between −1SD and +1SD) and high (total SRS > 1 SD). Each panel shows the relationship between age and the self-connection strength in the auditory cortex for the three groups. Contrary to what observed in low and mid SRS individuals, high symptom severity was associated with decreased A1 self-connection strength with age. Abbreviations: *SRS = Social responsiveness scale*.

## 4. Discussion

### 4.1 Reduced functional segregation of primary sensory regions in ASD

The establishment of a relative functional segregation between cortical and subcortical brain processing represents a crucial step in the development of distributed, functionally specialized cortico-cortical networks which characterize the architecture of the mature brain (K. Supekar, Musen, and Menon 2009). This architecture allows to maintain an equilibrium between local and distributed information processing, that is between functional segregation and functional integration (Caspers et al. 2013). The importance of such equilibrium is apparent during tasks which require both specialized processing and fast-paced integration of sensory stimuli, abstract concepts, and interoceptive information. Focused attention, language comprehension/production and social interaction are examples of such demanding everyday situations which are typically affected in autism.

In our analysis we observed that the functional segregation of primary somatosensory and auditory regions significantly increases with age in TD participants, while this effect is highly reduced in ASD participants. Probably the most interesting aspect of this result is the fact that in DCM, functional segregation - reflected in the parameter estimates of a brain region’s self-inhibition - is explicitly modelled as the contribution of GABA-ergic inhibitory projections to the sensitivity of each region (Bastos et al. 2012): a region that features low self-inhibition is more susceptible to be influenced by the activity of other regions (Zeidman, Jafarian, Corbin, et al. 2019). To our knowledge, the reduced segregation of primary sensory regions represents the first neuroimaging evidence of atypical development of the local brain circuitry mediated by inhibitory projections. At the same time it should be emphasized that while this association between intrinsic BOLD fluctuations and self inhibitory connections is supported by the explicit modelling of these GABA-ergic projections in the DCM model (Marreiros, Kiebel, and Friston 2008), the hypothesis of a direct link between BOLD activity and inhibitory projections remains tentative due to the intrinsic difficulties in determining to what extent the fMRI signal is determined by an (im)balance of excitatory and inhibitory activity (Logothetis 2008).

The investigation of inhibitory connections within and between cortical regions is of particular importance in autism research as many studies have proposed that this neuropsychiatric condition is associated with atypical development of intracortical inhibitory interneurons (Marín 2012; Le Magueresse and Monyer 2013; Ferguson and Gao 2018), resulting in an imbalance of local excitatory/inhibitory signalling (Rubenstein and Merzenich 2003; Belmonte et al. 2004). Importantly, local imbalance of excitatory/inhibitory projections affects not only local circuits, but also the development of long-range projections interconnecting large-scale networks (Menon 2013) due to delays in information transfer between distant regions and reduced synchrony in the activity of distant clusters of minicolumns (Belmonte 2004; Courchesne and Pierce 2005). This atypical connectional architecture is consistent with recent findings showing atypical development of whole brain functional segregation and integration in ASD, characterized by facilitated access of basic sensory information to higher-level cognitive processes (Hong et al. 2019; Martínez et al. 2020) and cross-talk between primary sensory and higher-order regions (Holiga et al. 2019; Martínez et al. 2020).

In the specific context of DCM, the lower intrinsic connectivity of the primary sensory regions reflects the smaller self-inhibition of these regions in ASD compared to TD participants. Since thalamic projections are predominantly glutamatergic (Salt 2002), this situation would allow an excessive influence of the excitatory afferents from the thalamus on cortical sensory processing. Such atypically high stimulus-driven circuitry is likely to decrease the efficiency of feedback projections modulating the thalamic afferents (Usrey and Sherman 2019; Briggs and Usrey 2008), and increase the saliency of the primary sensory information relayed to transmodal cortices and higher-order attentional processes. Similarly for the basal ganglia the excessive connectivity with primary sensory regions could hamper the development of effective fronto-striatal circuits which are involved in filtering out irrelevant or undesired stimuli. This possibility is supported by several reports of impaired sensorimotor gating in people with ASD (McAlonan et al. 2002; Perry et al. 2007; Madsen et al. 2014).

### 4.2 Persistent high influence of subcortical activity on cortical primary sensory regions in ASD

In a previous study we reported that ASD participants showed increased functional connectivity between subcortical and primary sensory regions (Cerliani et al. 2015). This effect could reflect an atypically enhanced (1) influence of subcortical activity on cortical sensory processing (bottom-up); (2) top-down modulation of subcortical activity; or (3) both. Since DCM is capable of separately modelling the directional influence between different regions, it represents the ideal modelling framework to disentangle which among these situations is more likely in ASD.

Given the prevalence of sensory symptoms in ASD, such as hyperreactivity to sensory stimulation and the presence of stereotyped and repetitive behaviour, as well as recent evidence of an increased flow of information from sensory to transmodal cortical regions (Hong et al. 2019), we specifically tested the hypothesis that the enhanced functional connectivity between subcortical and cortical regions in ASD would reflect an increased directional influence of subcortical activity on cortical processing in primary sensory regions. Our DCM analysis revealed that while in TD participants the influence of subcortical regions on primary sensory cortical regions decreases with brain maturation, this effect is largely not present in ASD participants. This situation could engender an excessive influence of unprocessed or undesired sensory information on cortico-cortical networks, overriding higher-order cognitive processes in determining the relevance of different cognitive representations to generate behavior.

While social and communicative deficits are central to the diagnosis of autism, the clinical literature has constantly remarked the importance of sensory symptoms in ASD, generally qualified as hyper- or hyporeactivity to sensory stimulation (Marco et al. 2011; Robertson and Baron-Cohen 2017; Cascio, Moore, and McGlone 2019). More specifically, sensory perception in ASD is characterized by enhanced perceptual processing and discrimination, which suggests a cognitive bias towards local over global features (Mottron et al. 2006; Minshew and Williams 2007; Robertson and Baron-Cohen 2017). At the same time, studies investigating basic measures of sensitivity in static sensory stimuli in autism failed to show higher thresholds for detection or discrimination in ASD than in TD. This recently led to the hypothesis that rather than a general bias towards local features, atypical sensory processing in ASD would particularly manifest in the slower dynamic integration of perceptual information over space and time, possibly due to inefficient or noisier sensory information processing in the brain (Robertson and Baron-Cohen 2017).

This hypothesis fits well in the framework of the underconnectivity/overconnectivity theory (Belmonte 2004; Just et al. 2004), according to which the global architecture of brain connections in autism could be characterized by inefficient long-range connections, supporting functional integration across different cognitive domains, coupled with an excess of local connections. The presence of local overconnectivity, consistent with decreased levels of GABA-ergic signalling and reduced minicolumnar size, would yield both a local and a distal effect on the development of brain connectivity. Locally, it would elicit indiscriminately high regional activation for any incoming sensory signal, thereby decreasing the selectivity of salient over irrelevant environmental stimuli (signal vs. noise) (Belmonte 2004). At the same time, increased local signalling would negatively impact the formation of long-distance projections, resulting in delayed and/or inefficient top-down modulatory projections, further decreasing the selectivity for salient environmental sensory information (Menon 2013).

On the basis of our results, subcortico-cortical overconnectivity in ASD could reflect excessive corticopetal flow of basic sensory information, resulting in decreased signal-to-noise ratio in primary cortical regions targeted by subcortical projections, due to the increased presence of irrelevant information (Belmonte et al. 2004). In turn, this would pose a challenge to attentional and higher-order cognitive networks in terms of fast-paced dynamic integration of the current sensory information. Such increased connectivity is apparent in ASD not only between subcortical and primary sensory regions, but also between the latter and transmodal regions which are directly connected to saliency and attentional networks, as suggested by recent findings of local overconnectivity at the transition between primary and transmodal sensory regions associated with a delayed transition of information to high-order regions in the fronto-parietal and default mode network (Hong et al. 2019).

Importantly, this evidence allows to reframe the idea of ‘local’ overconnectivity in terms of functional hierarchy of neural information processing (Mesulam 2000; Sepulcre et al. 2012; Margulies et al. 2016; Tian et al. 2020): while the hypothesis of long-range underconnectivity in ASD is consistent with many findings of decreased anatomical and functional connectivity between frontal and parietal brain regions (Just et al. 2012), evidence supporting local overconnectivity has only in part received a topographical localization (Jeffrey David Rudie and Dapretto 2013; Keown et al. 2013; Kaustubh Supekar et al. 2013). Indeed, while topographical proximity is generally a good predictor of anatomical or functional connectivity, the presence of distributed networks in the brain shows that distant regions can be more connected to each other than to regions which are topographically closer, but have different functional specialization. The same rationale applies also to the reciprocal connectivity between entire functional networks, which reflects the hierarchy of information processing in the brain (Margulies et al. 2016). Primary sensory and subcortical regions are topographically relatively distant, but they are monosynaptically connected with each other, and represent immediately subsequent steps in the information processing hierarchy. Our results showing higher connectivity between subcortical and primary sensory regions therefore supports the idea that local overconnectivity in ASD should be conceptualized, and investigated, both in terms of topographical and functional proximity.

## 5. Limitations

The use of solely cross-sectional samples represents a limitation to the current study which could hamper the interpretation of developmental effects and their interaction with the pathology. However, the ABIDE data mostly include only baseline measurement leaving no room for longitudinal modelling. Another limitation of the study is the presence of a statistical significant association between age and site, which prevents the possibility of effectively correcting the PEB model for the effect of scanning sites without removing variability in the DCM estimates that is potentially explained by age. Therefore, when adding dummy variables for each site, most of the model connections failed to reach statistical significance (Fig. S6, S7). However, as previously mentioned in the Methods, given the significant differences in mean age across sites (Fig. S5), the site confounds correlate with differences in Age, and therefore do not represent appropriate predictors for the unique variability associated with confounding differences between sites. For this reason, we presented the results using both with and without dummy variables for sites. In this context, we showed that the site-corrected and uncorrected models are very comparable in terms of directionality of results and the main findings (self-connection and bottom-up connection of A1) remain significant. Moreover, the ABIDE harmonized scan protocols and our centralized processing pipeline may have partially accounted for the effect of site.

## 6. Conclusion

In the present study we hypothesized that people with ASD would display a stronger flow of sensory information from subcortical nuclei to the cortex, which could explain the previously described increased functional connectivity between them. To test this hypothesis we modelled the bottom-up effective connectivity from basal ganglia and thalamus to the primary sensory regions in a relatively large group of participants (N=359). We found that (1) the influence of subcortical regions on primary visual and auditory cortices significantly decreased with age in TD, but not in ASD participants; (2) the functional segregation of somatosensory and auditory cortices from subcortical activity significantly increased with age only in TD participants, while this was the case only for the primary visual cortex in ASD participants. These results suggests that the recently detected increased and ectopic flow of information from primary sensory to higher order cortices in ASD (Hong et al. 2019; Holiga et al. 2019) originates already at the subcortical level, which is consistent with the decreased functional segregation of subcortical from cortical brain processes in ASD (Di Martino et al. 2011; Cerliani et al. 2015; Maximo and Kana 2019). The evidence of a specific directionality in this persistently high flow of sensory information from subcortical to cortical regions brings support to the idea that such hyperconnectivity could represent one of the brain mechanisms causing hyperreactivity to sensory stimuli in ASD.

## Acknowledgments

We thank Peter Zeidman for helping us with dynamic causal modelling analysis and interpretation, and for his comments on a previous version of the manuscript.

## Abbreviations

ABIDE: Autism Brain Imaging Data Exchange dataset
ASD: autism spectrum disorder
BMA: bayesian model average
BMR: bayesian model reduction
DCM: dynamic causal modelling
PEB: parametric empirical bayes
SRS: Social responsiveness scale
TD: typical development

