## Supplementary Information for "Atypical development of subcortico-cortical effective connectivity in autism"

#### Sample selection

**Image quality inclusion criteria**

(1) dataset includes a T1-weighted and RS-fmri scan with at least 180 time points (~5 min)
(2) absence of excessive artifacts (visual inspection on RS-fmri mean and stdev image over time - see Fig. S2)
(3) near-full brain coverage and succesful registration to T1-weighted image and MNI template

N = 545

↓

**Meeting basic phenotypical inclusion criteria**

Phenotypical inclusion criteria, based on the information provided by the Composite Phenotypic File available on the ABIDE website (Phenotypic_V1_0b.csv)
Sex=Male
Full IQ (FIQ) > 70
Age at scan available
Eye-status during scan available
Control participants (TD group) or participants with diagnosis of either Autism or Asperger's syndrome (ASD group)

N = 407
192 ASD
215 CON

↓

**Matching for Age, FIQ, Eye-status at scan**

Participants are matched by Age and Eye-status at scan in 8/8 sites, and by FIQ in 7/8 sites. The significant difference in FIQ in USM site (CON > ASD: p<.8E6) drives a significant difference in the FIQ in the entire group (CON > ASD: p<.004). To match participants by FIQ, a subsample of CON/ASD participants from the USM site with FIQ in the highest/lowest 20-th percentile are discarded.

N = 387
181 ASD
206 CON

↓

**Exclusion based on mean Framewise Displacement (mean FD)**

Mean FD is calculated according to Satterthwaite TD et al. (2012 Neuroimage) for each participant. Participants with mean FD higher than 0.34, corresponding to two standard deviations (2*0.1127) above the whole sample mean (0.1146) were excluded from further analysis. Specifically, 9 ASD and 5 CON participants were excluded.

N = 373
172 ASD
201 CON

↓

**Matching for mean FD**

Due to a residual difference in mean FD in the whole group (ASD > CON: p<.013), driven by participants in the NYU site (p<.043), CON/ASD participants from the NYU site with mean FD in the lowest/highest 5-th percentile are discarded (6 ASD and 8 CON). After this exclusion, remaining participants are matched by Age (p=.39), FIQ (p=.35) Eye-status at scan (p=.81) and mean FD (p=.09). In subsequent analyses, all these parameters, plus a variable coding for the acquisition site, are used as covariates of no interest.

N = 359
166 ASD
193 CON

**Figure S1.** Pipeline used to derive the final sample of ASD and TD participants data used for the present study, as well as for our previous functional network connectivity study (Cerliani et al. 2015).

### Preprocessing and selection of the regions of interest based on a previous functional connectivity study

All the preprocessing steps and the subsequent independent component analysis (ICA) - which was used to define the seed regions of interest for the present effective connectivity study - are described in detail in a previous study (Cerliani et al. 2015).

To summarize, we used FSL to preprocess the resting-state fMRI images including brain extraction, linear motion correction, slice timing correction, spatial smoothing, and linear registration to the T1-weighted anatomical scan. To define the regions of interest which would then be used to extract the subject-level characteristic time courses to feed into the effective connectivity analysis, we used the results of a previous study of ours (Cerliani et al. 2015) which employed independent component analysis (ICA) on the same subset of 359 participants from the ABIDE database (Di Martino et al. 2013). This choice was motivated by the fact that the results of that study provided the evidence of increased subcortico-cortical connectivity that represents the ground for our current hypotheses about the atypical directional influence of subcortical regions onto primary sensory regions in ASD which is currently tested using dynamic causal modelling (Friston et al. 2014).

ICA was carried out using FSL Melodic (Beckmann and Smith 2004). The spatially independent components extracted by ICA represent the regions of interest used in the previous as well as in the present study.

To estimate robust spatial independent components, temporally concatenated melodic ICA was carried out 25 times on subsets of 112 participants randomly chosen from the dataset, and the final spatial components were obtained by a meta-ICA on the results of the first-level ICAs. Out of the resulting 52 automatically estimated meata-ICA components, we selected 19 of them mostly located in the gray matter, featuring high reproducibility across subsets, high resemblance to functional networks recruited by task-based fMRI experiments, temporal frequency spectrum in the low-frequency range of resting-state networks.

After thresholding the 19 spatial components at $Z > 3$ (Z values estimated by meta-ICA), their characteristic time course was extracted via dual regression (Nickerson et al. 2017), bandpass filtered (0.009-0.08 Hz), and used to estimate functional connectivity between networks within each participant, with the aim of subsequently comparing these functional connectivity estimates between groups after regressing out the effect of age, IQ, eye status (open/closed) at scan, motion (mean framewise displacement) and site of acquisition (one dummy variable for each site - 1 to prevent rank deficiency)

Inference on the difference in functional connectivity between ASD and TD participants was carried out using permutation testing (Nichols and Holmes 2002) with 20,000 permutations for each cell of the 19x19 functional connectivity matrix. The resulting p values (from a two-tailed test) were corrected for multiple comparison using false discovery rate (Genovese, Lazar, and Nichols 2002) (q(FDR)= 0.05) considering all the p values in the upper triangular connectivity matrix, that is 19x18/2 p values (identical to the values in the lower triangular as the functional connectivity matrix is symmetric). This resulted in a final corrected p threshold of 0.001. This analysis revealed that functional connectivity between one subcortical network - encompassing the basal ganglia and thalamus - and four primary sensory cortical networks - ventral and dorsal somatosensory, visual and auditory - was significantly higher in ASD than in TD. No other group-differences were significant after correction for multiple comparisons (Figure S2). These 5 networks were used in the present study as regions of interest (ROI) for dynamic causal modelling (Friston et al. 2014).

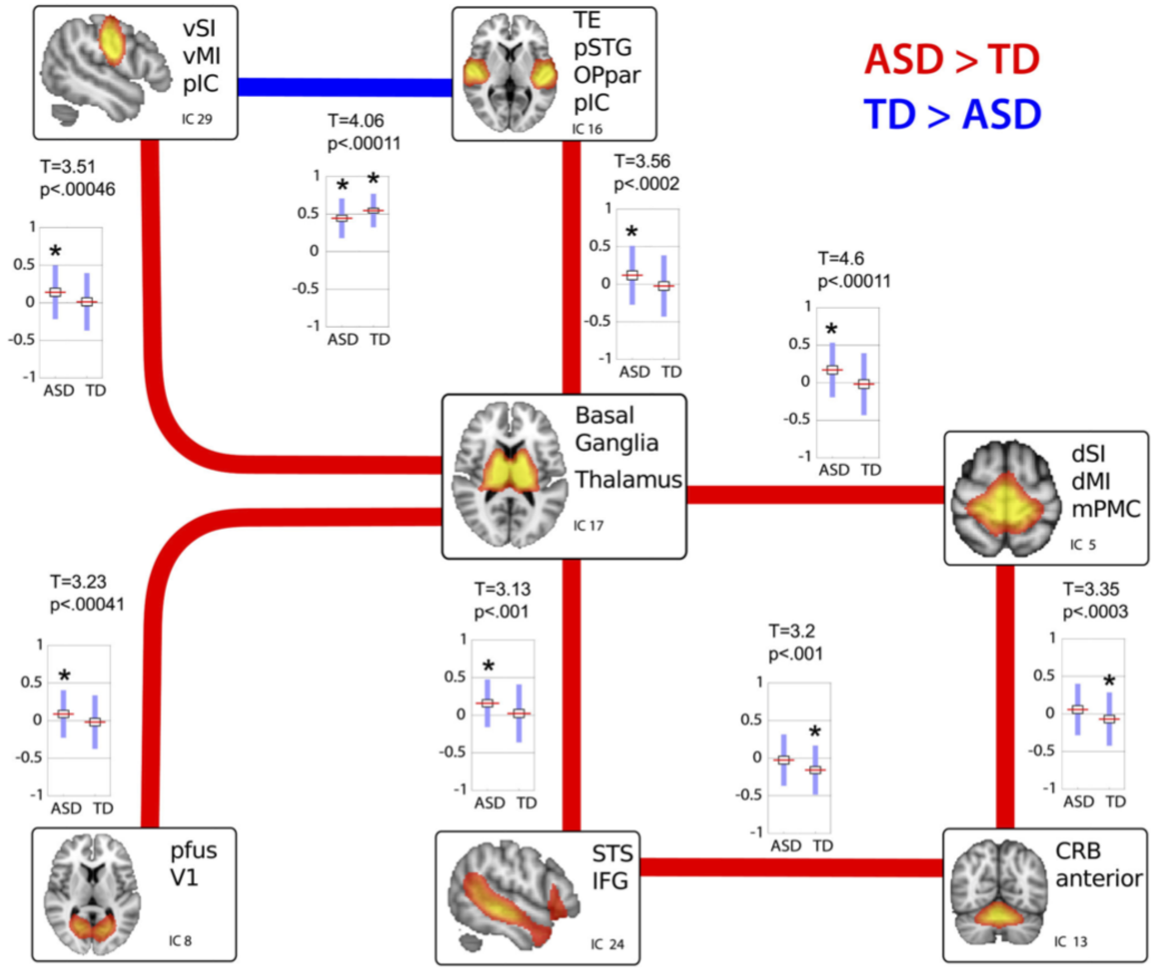


**Figure S2** (adapted from Fig. 2 in (Cerliani et al. 2015), available at https://pubmed.ncbi.nlm.nih.gov/26061743/): Group differences in functional network connectivity strength are shown as lines (red indicates increased functional network connectivity in autism spectrum disorder [ASD] with respect to typically developing [TD] participants, and blue indicates the reverse situation) together with boxplots of the Pearson product moment correlation values per each group. Boxplots report the mean, SEM (blue rectangle around the mean), and SD (whiskers) of group-level functional network connectivity values. Results were obtained by comparing the between-network functional connectivity of 166 ASD group and 193 TD group participants using nonparametric permutation testing (20,000 permutations) and correcting the final results with q[FDR] = 0.05, leading to a final threshold of $P < .001$. Converting the correlation scores to z scores using Fisher r to z transformation yielded almost identical results (eFigure 5 in the original study). The expanded independent component (IC) abbreviations are listed in Table 2. FDR indicates false discovery rate.

### Spectral dynamic causal modelling

To identify the effective connectivity between the 5 ICA-based spatial maps (IC17 - basal ganglia and thalamus; IC5 dorsal somatosensory; IC29 - ventral somatosensory; IC8 - primary visual; IC16 - primary auditory) we employed spectral dynamic causal modelling (spDCM) (Friston et al. 2014). This framework is of particular interest in modelling resting-state fMRI time-series due to the parameterization of endogenous fluctuations by characterizing them in terms of their cross-spectral densities. Moreover, compared to the previous version of DCM for resting state (i.e. stochastic DCM) this eliminates the need of estimating the neural fluctuations speeding up and simplifying model inversion (Razi et al. 2015).

DCM was carried out using SPM 12. For each subject, the full model included all the bottom up (subcortico-cortical) and top down (cortico-subcortical) connections between our regions of interest, and the inhibitory self-connections within each region, by setting the prior variance to 1 allowing them to be informed by the data. Since direct connections between primary sensory cortices are anatomically implausible (Mesulam 2000), cortico-cortical connections were not modelled, setting the prior variance to zero, reducing the number of parameters to be estimated.

The model inversion for each subject provided the estimation of the connection strength parameters which best explained the observed data, resulting in the expected value and covariance (uncertainty) for each connection, as well as the free energy approximation to log model evidence, indicating the quality of each model in terms of accuracy and complexity.

### Parametric empirical Bayes

We subsequently tested differences in parameter strength due to between subject variability in the diagnosis, the age and the interaction between these two. To do so, we used a recent implementation of SPM to model group-level connectivity in the context of DCM, known as parametric empirical Bayes (Friston et al. 2016). This can be considered as a hierarchical Bayesian second-level general linear model that models how subject measures (individual connection strengths) relate to the group mean and other group-level variables. Unlike previous approaches to investigate group differences between DCM parameters which rested on classical inference approaches on individual expected values, this routine has the advantage of taking into consideration the full posterior density from the first level DCM, including both the estimated strength of each connection and the uncertainty (covariance), to inform the second level results (Zhou et al. 2018).

We therefore created a general linear model with 4 mean-centered variables of interest, i.e. the mean, the group, the age and the interaction between age and group, as the multiplication of age by group. We also included one nuisance variable with the mean framewise displacement (FD) for each subject in order to exclude potential effects of movement, allowing our PEB model to express the influence of our between-subject variables (each column of the design matrix) on the within-subject measures (parameter strength) as well as estimating the between-subject variability.

### Bayesian model reduction

After having inverted the individual full model and the group PEB model we used Bayesian model reduction and comparison to evaluate differences of effective connectivity across subjects. With this routine, several nested (reduced) PEB models are specified assuming one or more connections from the full model to be selectively switched off, and are then rapidly estimated deriving the model evidence directly from the full model. The Bayesian model average is subsequently used to estimate a weighted average of the parameter strength using the log evidence of the nested models (Friston et al. 2016).

Because we were interested in specific bottom-up or top-down connections, we constrained the Bayesian model reduction to 60 templates by selectively switching off just bottom-up or top-down connections.

To sum up, we first fitted a full model on the individual data obtaining the estimation of the parameters (connections) for each subject, we then brought this to the second (group) level using parametric empirical Bayes and finally we performed Bayesian model reduction and averaging. A schematic representation of the analysis pipeline is shown in figure S2.

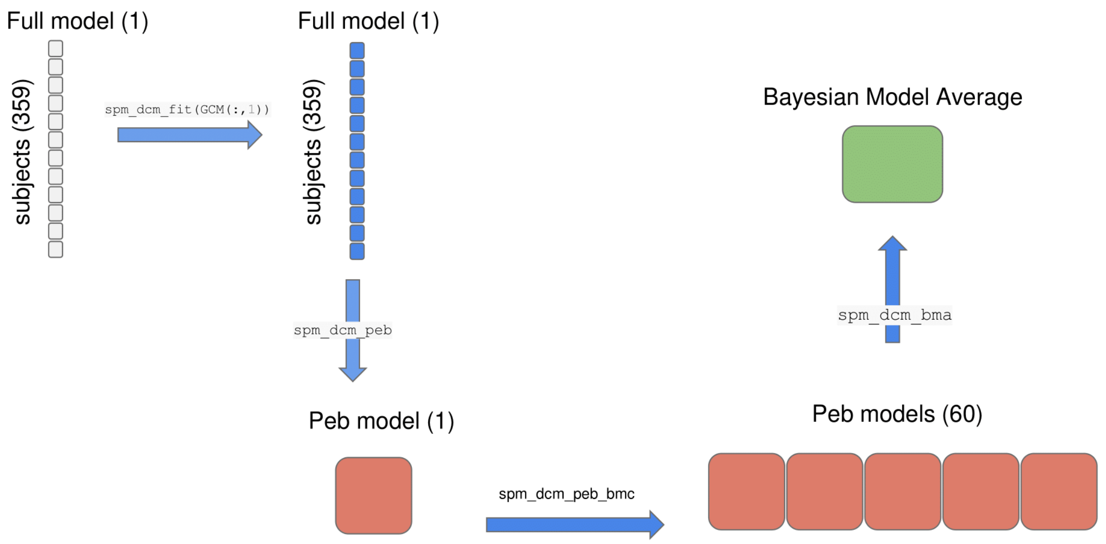


**Figure S3 A brief overview of the DCM pipeline** (adapted from (Friston et al. 2016)**.** The unfilled squares correspond to the specified models while the filled one to fitted models. Each column of the array corresponds to a particular model (with the full model on the first column). Each row corresponds to a particular subject. The figure illustrates how the analysis has been carried out. Firstly, we inverted the full model to the data obtaining the posterior probability and the parameter strengths. We then created a single PEB model (spm_dcm_peb) from the single subject full models and compared this with reduced PEB models (spm_dcm_peb_bmc). Finally we examined the BMA over the 60 PEB models.

### Supplementary Results

#### Participants' Age

Distribution of age across the whole sample included in the analysis is reported in Figure S4.

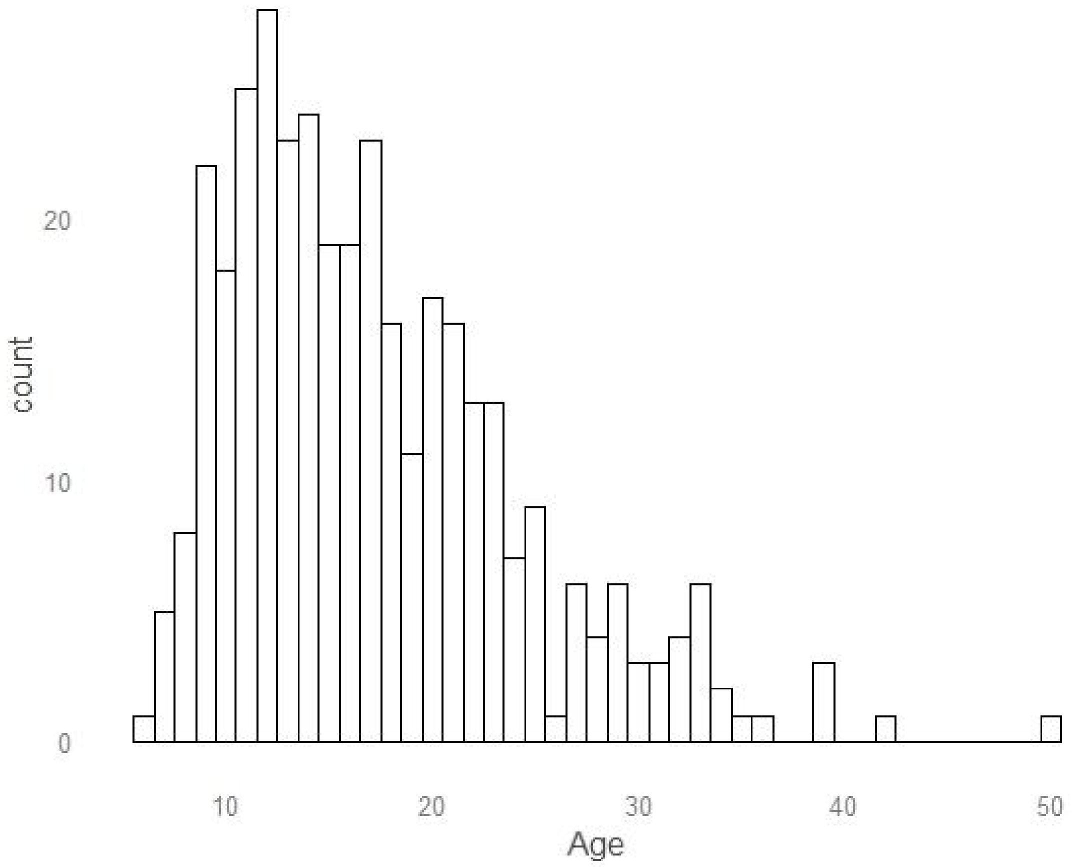


**Figure S4. Age distribution across the whole sample.**

As reported in the manuscript, using analysis of variance (ANOVA) we found differences in the mean age across sites significant ($F(7,351) = 22.07$, $p < 0.001$). Figure S3 illustrates age distribution across the 8 sites.

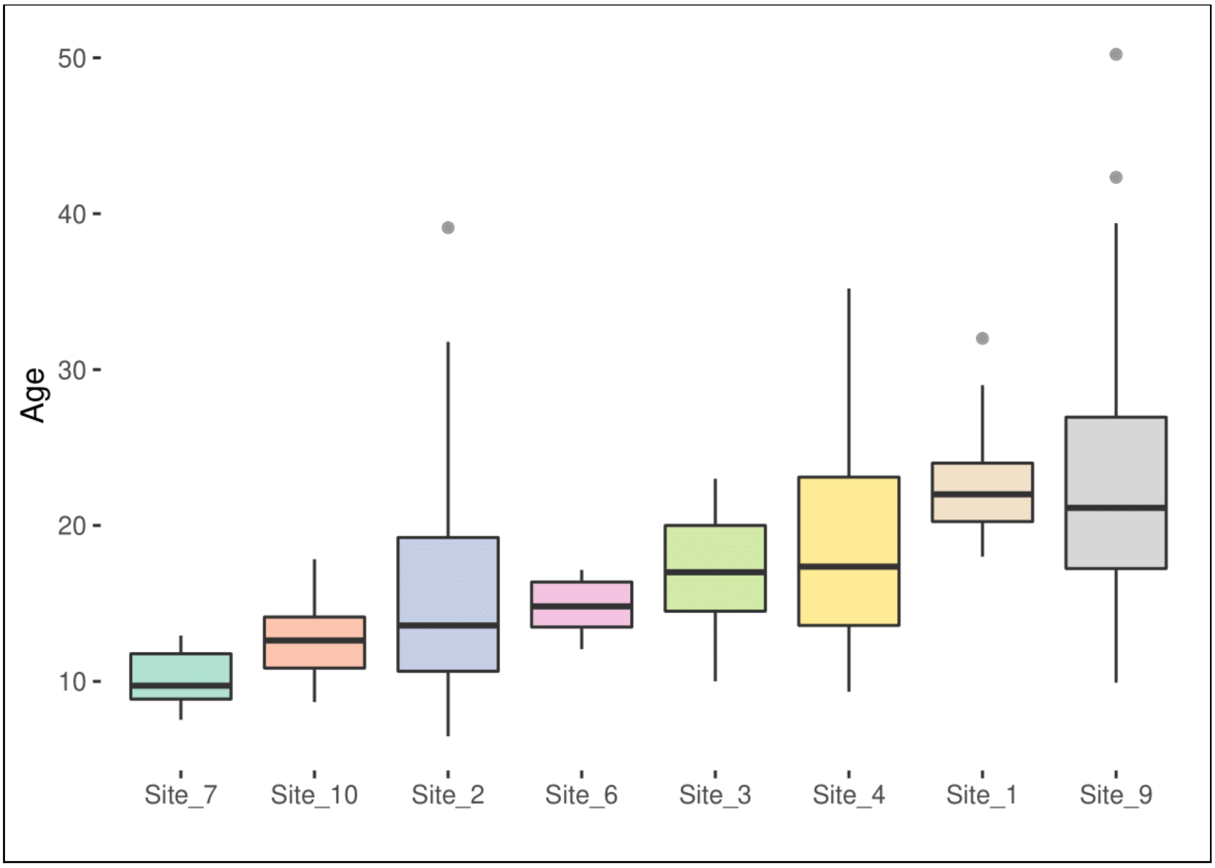


**Figure S5. Mean age differences across sites.** Differences in age between sites, ordered from the lowest to the highest age per site.

#### Effect of age on network connectivity in the whole group

We found a main effect of age on DCM connections. As discussed in the text, intrinsic self-inhibition of both primary sensory and subcortical networks increased with age. This effect likely reflects the consolidation of the intrinsic circuitry within a functionally specialized network, and the progressive functional segregation between subcortical and cortical networks which occurs during development. Moreover, age was generally associated with a decrease in bottom-up connections, reaching primary sensory cortices from subcortical nuclei and an increase of top-down feedback cortical afferents to the basal ganglia and thalamus. Figure S6 shows SPM graphical output of this relationship and the effect of correcting for site. However, as mentioned in the main text and shown in the previous section, site was related with age and created therefore multicollinearity in the PEB model.

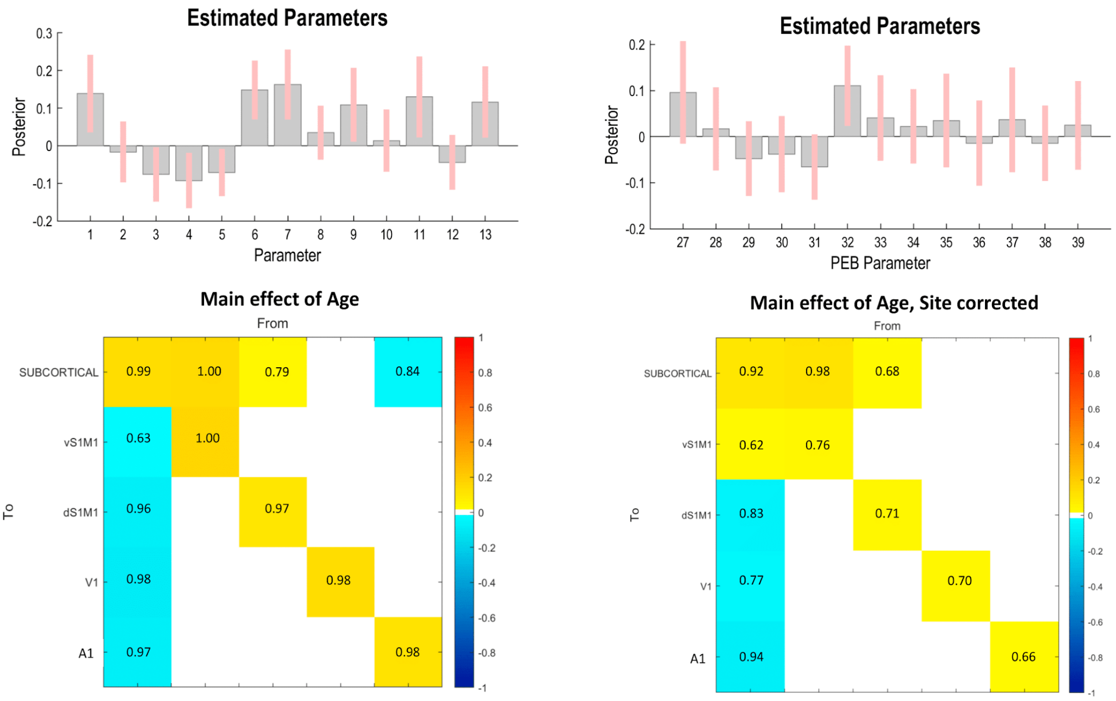


**Figure S6. Main effect of Age.** SPM graphical output when displaying the influence age on DCM parameters. Left: output of the analysis run without correcting for site with barplot (*top*) showing effect sizes (gray bars) and 90% confidence intervals (pink lines), and the effective connectivity matrix (*bottom*) showing the main effect of age on intrinsic connectivity for each region (values on the diagonal), bottom-up (values in the first column) and top-down (first row) connections. Colors show estimated effect in Hz while annotated numbers show posterior probabilities for each connection. *Right*: results of the same analysis after site correction.

### Effect of the interaction between age and group on network connectivity

Our main result was found looking at the effect of the interaction between age and group on DCM parameters. In brief, while TD showed a typical development of subcortico-cortical connections, with a weaker influence of basal ganglia and thalamus on primary sensory cortices, in ASD participants we found an attenuated functional segregation of sensory networks. Here we report the SPM graphical output of this analysis and the effect of adding site as a covariate in the PEB model (Figure S5).

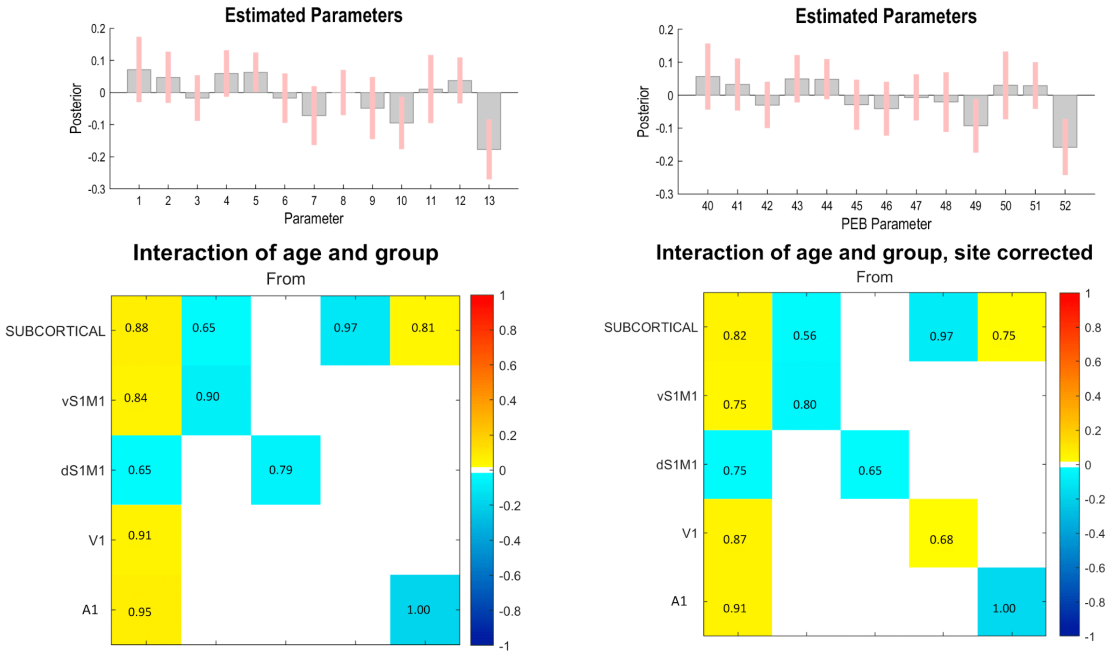


**Figure S7. Interaction between age and group.** SPM graphical output when displaying the influence of the interaction between age and group on DCM parameters. Left: output of the analysis run without correcting for site with barplot (*top*) showing effect sizes (gray bars) and 90% confidence intervals (pink lines), and the effective connectivity matrix (*bottom*) showing the effect of the interaction term on intrinsic connectivity for each region (values on the diagonal), bottom-up (values in the first column) and top-down (first row) connections. Colors show estimated effect in Hz while annotated numbers show posterior probabilities for each connection. *Right*: results of the same analysis after site correction.

### Association with Social Behaviour

**Table S1. Linear mixed models investigating association of SRS, age and their interaction on estimated DCM parameter strengths.**

| *Connection* | *Contrast* | *Estimates* | *Confidence Interval* | *P-Value* | *P-Value (FDR corrected)* |
|---|---|---|---|---|---|
| vS1 self connection | | | | | |
| | Age | -0.07 | -0.46-0.33 | 0.74 | |
| | SRS | 0.001 | -0.05-0.05 | 0.69 | |
| | Age*SRS | 0.001 | -0.01-0.01 | 0.91 | 0.916 |
| A1 self connection | | | | | |
| | Age | 0.05 | -0.33–0.42 | 0.81 | |

| | | | | | |
|---|---|---|---|---|---|
| | SRS | -0.01 | -0.01-0.01 | 0.81 | |
| | Age*SRS | -0.01 | -0.01-0.01 | **0.013*** | **0.053*** |
| Subcortical to V1 | | | | | |
| | Age | 0.04 | -0.32-0.4 | 0.84 | |
| | SRS | -0.01 | -0.01-0.01 | 0.17 | |
| | Age*SRS | -0.01 | -0.01-0.01 | 0.71 | 0.916 |
| Subcortical to A1 | | | | | |
| | Age | -0.13 | -0.37-0.11 | 0.29 | |
| | SRS | -0.01 | -0.01-0.01 | 0.19 | |
| | Age*SRS | -0.01 | -0.01-0.01 | 0.52 | 0.916 |

Nickerson, Lisa D., Stephen M. Smith, Döst Öngür, and Christian F. Beckmann. 2017. "Using Dual Regression to Investigate Network Shape and Amplitude in Functional Connectivity Analyses." *Frontiers in Neuroscience* 11 (March): 115.

Razi, Adeel, Joshua Kahan, Geraint Rees, and Karl J. Friston. 2015. "Construct Validation of a DCM for Resting State fMRI." *NeuroImage* 106 (February): 1–14.

Zhou, Yuan, Peter Zeidman, Shihao Wu, Adeel Razi, Cheng Chen, Liuqing Yang, Jilin Zou, Gaohua Wang, Huiling Wang, and Karl J. Friston. 2018. "Altered Intrinsic and Extrinsic Connectivity in

Schizophrenia." *NeuroImage. Clinical* 17: 704–16.
